# The invisibility cloak: Chitin binding protein of *Verticillium nonalfalfae* disguises fungus from plant chitinases

**DOI:** 10.1101/462499

**Authors:** Helena Volk, Kristina Marton, Marko Flajšman, Sebastjan Radišek, Ingo Hein, Črtomir Podlipnik, Branka Javornik, Sabina Berne

**Affiliations:** Department of Agronomy, Biotechnical Faculty, University of Ljubljana, Jamnikarjeva 101, SI-1000 Ljubljana, Slovenia; Slovenian Institute of Hop Research and Brewing, Cesta Žalskega tabora 2, SI-3310 Žalec, Slovenia; The James Hutton Institute, Invergowrie, Dundee DD2 5DA, Scotland, United Kingdom; The University of Dundee, School of Life Sciences, Division of Plant Sciences at the JHI, Invergowrie, Dundee DD2 5DA, Scotland, United Kingdom; Department of Chemistry and Biochemistry, Faculty of Chemistry and Chemical Technology, University of Ljubljana, Večna pot 113, SI-1000 Ljubljana, Slovenia

**Author notes:** Corresponding author: S. Berne. Nucleotide sequence data is available under accession numbers MH325205 for *VnaChtBP*, and MH325206 for *VaChtBP*.

## Abstract

During fungal infections, plant cells secrete chitinases that digest chitin in the fungal cell walls. The recognition of released chitin oligomers via lysin motif (LysM)-containing immune receptors results in the activation of defence signalling pathways. We report here that *Verticillium nonalfalfae*, a hemibiotrophic xylem-invading fungus, prevents this recognition process by secreting a CBM18 (carbohydrate binding motif 18)-chitin binding protein, VnaChtBP, which is transcriptionally activated specifically during the parasitic life stages. VnaChtBP is encoded by the *Vna8.213* gene which is highly conserved within the species, suggesting high evolutionary stability and importance for the fungal lifestyle. In a pathogenicity assay, however, *Vna8.213* knockout mutants exhibit wilting symptoms similar to the wild type fungus, suggesting that *Vna8.213* activity is functionally redundant during fungal infection of hop. In binding assay, recombinant VnaChtBP binds chitin and chitin oligomers *in vitro* with submicromolar affinity and protects fungal hyphae from degradation by plant chitinases. Using a yeast-two-hybrid assay, homology modelling and molecular docking, we demonstrated that VnaChtBP forms dimers in the absence of ligands and that this interaction is stabilized by the binding of chitin hexamers with a similar preference in the two binding sites. Our data suggest that, in addition to chitin binding LysM (CBM50) and Avr4 (CBM14) fungal effectors, structurally unrelated CBM18 effectors have convergently evolved to prevent hydrolysis of the fungal cell wall against plant chitinases.

## Introductory statements

Plant defense against pathogenic organisms relies on innate immunity, which is triggered by recognition of pathogen-derived or endogenous danger signals by plant receptors, described as pattern-triggered immunity (PTI) and effector-triggered immunity (ETI) (Jones and Dangl 2006; Dodds and Rathjen 2010). PTI, as a first line of defense, is activated by host cell surface-localized pattern recognition receptors (PRRs) sensing pathogen- and danger-associated molecular patterns (PAMPs and DAMPs, respectively) (Boller and Felix 2009; Böhm et al. 2014), which can be suppressed by pathogen virulence effectors (Dou and Zhou 2012). Host recognition of pathogen effectors by cytoplasmic nucleotide-binding domain leucine-rich repeat receptors (NLRs) leads to a second line of defense, ETI (Jones and Dangl 2006; Spoel and Dong 2012).

Pattern recognition receptors, which are either receptor-like kinases (RLKs) or receptor-like proteins (RLPs) that function in conjunction with RLKs, sense PAMPs or DAMPs and transduce downstream signaling to trigger PTI responses. Early PTI responses include the rapid generation of reactive oxygen species, activation of ion channels and mitogen-activated protein kinases. In turn, this leads to the expression of defense related genes, leading to an accumulation of antimicrobial compounds such as enzymes, which damage pathogen structures, inhibitors of pathogen enzymes and other antimicrobial molecules (Boller and Felix 2009; Dodds and Rathjen 2010; Macho and Zipfel 2014).

PAMPs, released during infection, are conserved molecular patterns characteristic of different pathogen classes (Ranf 2017). In fungi, chitin, in addition to beta-glucan and xylanase, is a well-studied PAMP that activates the host defense response. Chitin (a polymer of β-1,4-linked N-acetylglucosamine; (GlcNAc)_n_), is a major and highly conserved component of fungal cell walls and can be degraded to chitin oligosaccharides by plant apoplastic chitinases. The generated chitin fragments are recognized by a chitin perception system and subsequently activate PTI (Shibuya and Minami 2001; Sanchez-Vallet et al. 2014; Shinya et al. 2015).

Major chitin sensing PRRs, RLKs and RLPs belonging to the LysM domain family, are well studied in *Arabidopsis* and rice (Gust et al. 2012; Ranf 2017). *Arabidopsis* LysM-RLK AtCERK1 (chitin elicitor receptor kinase1) binds N-acetylated chitin fragments with three LysM motifs and, through homodimer formation, mediates chitin-inducible plant defenses (Miya et al. 2007; Liu et al. 2012). Cao et al. (2014) later identified another LysM-RLK in *Arabidopsis*, AtLYK5, which binds chitin at higher affinity than AtCERK1. The authors propose that AtLYK5 functions as the major chitin receptor, which recruits AtCERK1 to form a chitin inducible receptor complex. In rice, two receptors are involved in chitin triggered immunity. LysM-RLP OsCEBiP (chitin elicitor binding protein) binds N-acetylated chitin fragments, which initiates receptor homodimerization and further heterodimerization with OsCERK1. This heterotetramer formation triggers chitin induced PTI (Hayafune et al. 2014).

To overcome chitin-triggered immunity, successful pathogens have evolved various strategies, including alteration of the composition and structure of cell walls, modification of carbohydrate chains and secretion of effector proteins to prevent hydrolysis of the fungal cell wall or the release and recognition of chitin oligosaccharides (Sanchez-Vallet et al. 2014).

A well-described strategy of fungal cell wall protection against host chitinases is that of the tomato leaf mold fungus *Cladosporium fulvum*, which secretes chitin-binding protein Avr4 during infection. Avr4 effector binds with its carbohydrate-binding module family 14 (CBM14) to the fungal cell wall chitin and thus shields fungal hyphae against degradation by chitinases (van den Burg et al. 2006; van Esse et al. 2007). There is evidence for a similar protection of cell wall chitin in a phylogenetically closely related species of the Dothideomycete fungi class harboring homologs of Avr4 (Stergiopoulos et al. 2010). Protection of fungal hyphae against hydrolysis by chitinases has also been shown for Mg1LysM and Mg3LysM of *Zymoseptoria tritici* (formerly *M. graminicola*) (Marshall et al. 2011). Furthermore, Vd2LysM from *Verticillium dahliae* (Kombrink et al. 2017) belongs to LysM fungal effectors which are known to bind chitin oligomers. The first LysM effector, Ecp6, was found in the tomato pathogen *C. fulvum* and its characterization provided evidence that Ecp6 specifically and with high affinity binds chitin oligosaccharides. This competition with receptors subsequently disrupts chitin recognition by host receptors and suppresses the chitin-triggered immune response (Bolton et al. 2008; de Jonge et al. 2010; Sánchez-Vallet et al. 2013). Genomes contain several genes for LysM effectors and those highly expressed during infection have been characterized in fungal pathogens, including *Z. tritici* (Marshall et al. 2011), *Magnaporthe oryzae* (Mentlak et al. 2012), *Colletotrichum higginsianum* (Takahara et al. 2016) and *V. dahliae* (Kombrink et al. 2017). These studies demonstrate the involvement of LysM effectors in pathogen virulence by scavenging chitin oligomers to prevent recognition by the host chitin receptors, thus suppressing the chitin-triggered immunity.

The question arises if there are other molecules/systems/complexes apart from Avr4 (CBM14) and LysM (CBM50) effectors that can interfere with plant chitin perception and activation of PTI. We have been studying the *Verticillium nonalfalfae* – hop (*Humulus lupulus* L.) pathosystem. In an early comparative transcriptomic study of compatible and incompatible interactions (Cregeen et al. 2015), an *in planta* expressed *V. nonalfalfae* lectin gene was detected. A preliminary study showed that this *V. nonalfalfae* lectin contains putative carbohydrate-binding module family 18, CBM18 (Wright et al. 1991) domains. CBM18 is a chitin-binding domain involved in recognition of chitin oligomers and typically found in fungal and plant proteins in one or more copies (Lerner and Raikhel 1992). We report here on the characterization of *V. nonalfalfae* lectin with six CBM18 domains and show that it is a novel effector in plant fungal pathogens. CBM18 binds chitin and protects hyphae of *Trichoderma viride* from hop chitinases in an *in vitro* protection assay.

## Results

### The majority of CBM18 module containing proteins of *V. nonalfalfae* are expressed *in planta*

The *Vna8.213* gene, encoding a putative pathogen CBM18-containing chitin binding protein (VnaChtBP), has previously been identified as a differentially expressed transcript during compatible and incompatible interactions of *V. nonalfalfae* and hop (Cregeen et al. 2015). Surveying the *V. nonalfalfae* genome (Jakše et al. 2018) uncovered ten additional genes that encode for proteins with at least one CBM18 module (Fig. 1). These genes were grouped into four categories according to their domain architecture: Lectin-like proteins (Fig. 1A), Chitinases (Fig. 1B), Chitin deacetlyases (Fig. 1C) and Xyloglucan endotransglucosylase (Fig. 1D). The size of these proteins ranged between 349 and 1,696 amino acids (Vna6.1 and Vna1.668, respectively) and they harbored between one to ten CBM18 modules. Amongst these genes, ten are differentially expressed *in planta* (Fig. 1E) (Marton et al. 2018) and five (Vna2.980, Vna6.6, Vna8.213, Vna9.506 and Vna9.510) were predicted to be classically secreted proteins with N-terminal signal peptides. Amongst the chitinases (Fig. 1B), transcripts of *Vna3.655* and Vna9.506 were detected exclusively in susceptible hop, *Vna1.668* transcripts were found expressed in the roots of both resistant (‘Wye Target’) and susceptible (‘Celeia’) hop varieties, while transcripts of *Vna2.980* and *Vna9.510* were barely detectable. Interestingly, only one chitin deacetylase gene (VnaUn.355) was expressed during infection, and it showed preferential induction in the roots of both hop varieties. Such an expression profile was also evident for transcripts of *Vna6.6* belonging to xyloglucan endotransglucosylase. The highest expression was observed for *Vna8.213* transcripts, in particular at the late stages of infection of susceptible hop. Interestingly, *Vna1.667* gene-encoding lectin-like protein, containing 10 CBM18 modules, was barely expressed in the roots of susceptible hop during the early infection stages.

**Fig. 1.**
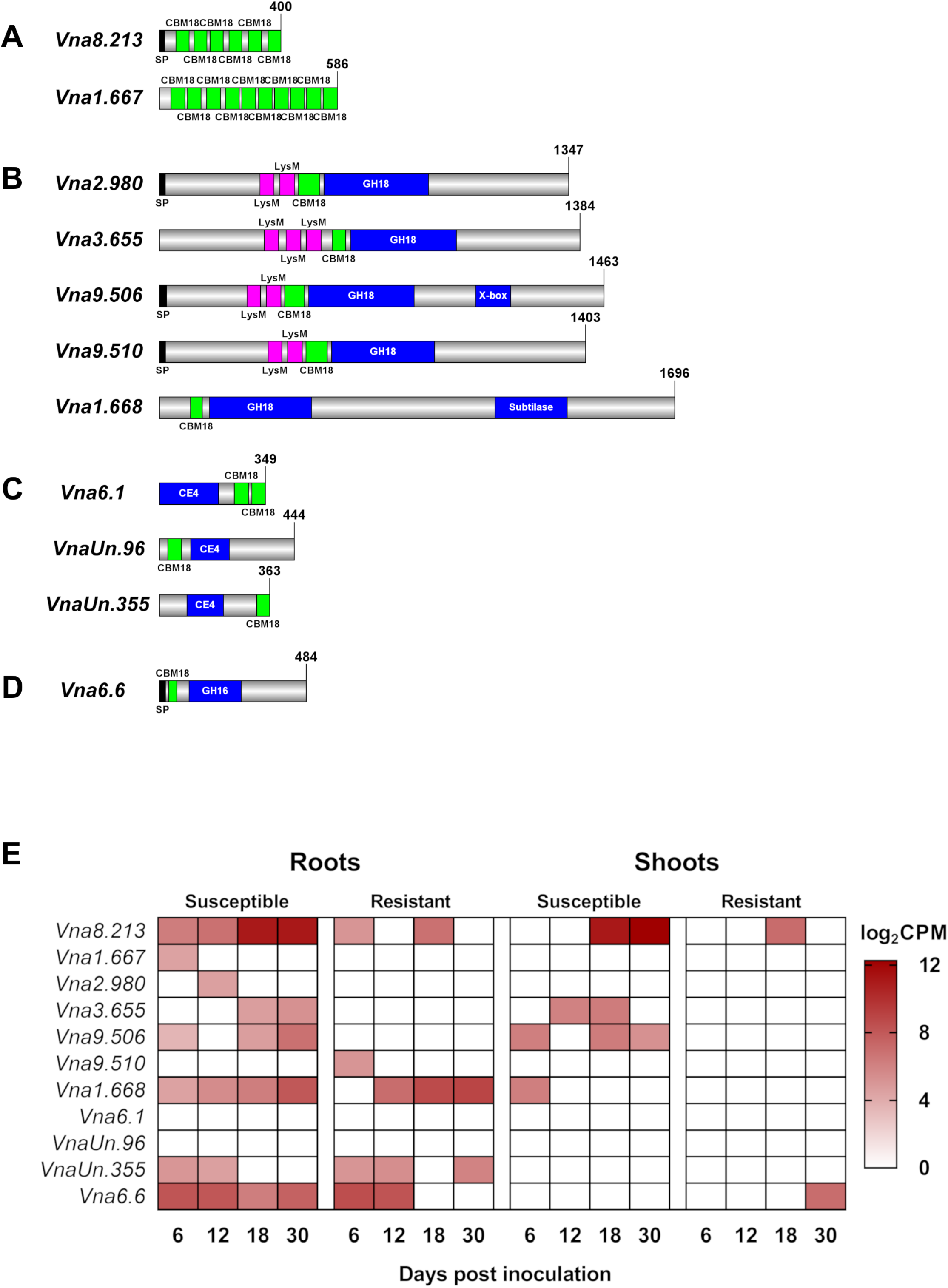
Domain architecture (A-D) and gene expression (E) of CBM18-containing proteins identified in *Verticillium nonalfalfae*. Protein organization was determined by querying protein sequences against CATH-Gene3D (Dawson et al. 2017) using the FunFHMMer web server and presented by IBS software (Liu et al. 2015). Proteins were classified into four groups: Lectin-like proteins (A), Chitinases (B), Chitin deacetlyases (C) and Xyloglucan endotransglucosylase (D). Gene expression is presented as a heatmap of log_2_CPM values determined by RNA sequencing of infected hop (Progar et al. 2017).

To confirm the expression patterns of *Vna8.213* measured by RNA-Seq, detailed gene expression profiling of root and shoot samples from susceptible and resistant hop varieties was performed using RT-qPCR at 6, 12 and 18 days post-inoculation (dpi) with *V. nonalfalfae* (Fig. 2). Gene expression of *Vna8.213*, from here on designated as *VnaChtBP*, increased with time, reaching the highest abundance in stems of susceptible hop at 18 dpi. The overall *VnaChtBP* expression in resistant hop was at a much lower level than in the susceptible variety and peaked at 12 dpi in stems. These results indicate that *VnaChtBP* expression is induced *in planta* and its transcript abundance in susceptible hop increases with progression of fungal colonization.

**Fig. 2.**
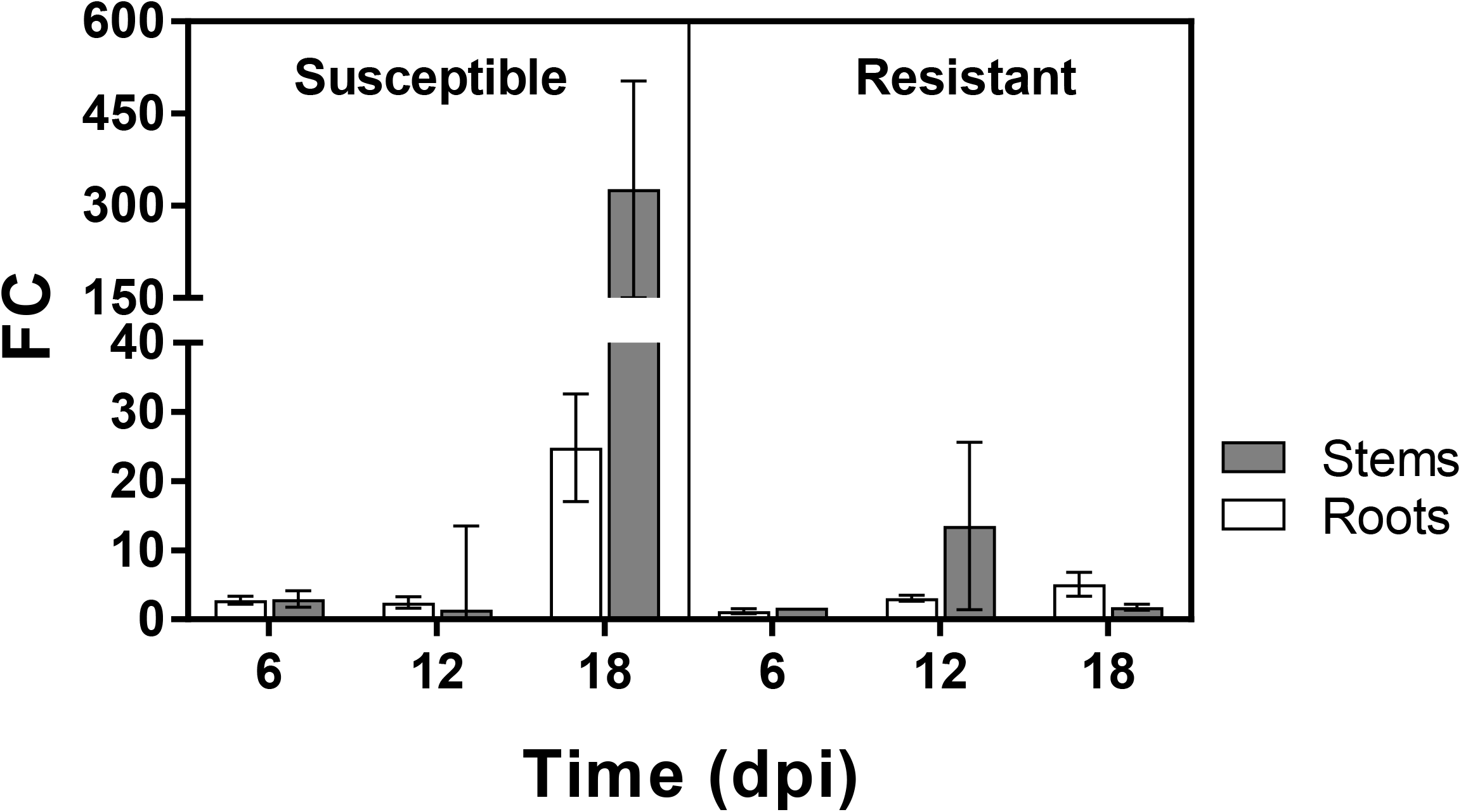
*VnaChtBP*, a gene encoding the CBM18 chitin binding protein of *Verticillium nonalfalfae*, is highly expressed in stems of susceptible hop at the late stages of infection. The gene expression of *VnaChtBP* was quantified by RT-qPCR using a cDNA prepared from the roots and shoots of infected susceptible (‘Celeia’) and resistant (‘Wye Target’) hop plants (n = 5) at 6, 12 and 18 dpi and the expression levels were normalised relative to the expression of the gene in ½ liquid Czapek-Dox medium using topoisomerase (*VnaUn.148*) and splicing factor 3a2 (*Vna8.801*) as housekeeping genes (Marton et al. 2018). FC, fold change; dpi, days post inoculation.

### Sequence conservation suggests evolutionary stability of *VnaChtBP*

To investigate the presence and sequence variation of *VnaChtBP* in 28 *V. nonalfalfae* isolates (Suppl. Table S1), PCR amplification and Sanger sequencing of cloned genes was performed. The *VnaChtBP* gene was present in all analyzed isolates and displayed no sequence polymorphisms. This is consistent with purifying selection and suggests evolutionary stability of the gene as well as an important role in the fungal lifestyle.

Among all sequences deposited at NCBI, VnaChtBP shared the highest protein identity with a lectin from *V. alfalfae* (97%; an alfalfa isolate VaMs.102), followed by *V. dahliae* lectin-B (80%; a lettuce isolate VdLs.17), two *V. dahliae* hypothetical proteins, Vd0004_g7043 and Vd0001_g7025 (80% and 79%; strawberry isolates 12161 and 12158), and a hypothetical protein BN1708_012400 from *V. longisporum* (78%; a rapeseed isolate VL1) (Suppl. Table S2). Additional homologs (Suppl. File S1), but with lower identity (48-39%), were identified in fungi amongst Sordariomyceta (40) and Dotideomyceta (3), and in fungi *Incertae sedis* amongst Neocallimastigomycetes (5) and Chytridiomycetes (2).

Due to the high sequence similarity shared between VnaChtBP and *V. alfalfae* VaMs.102 lectin, PCR screening and Sanger sequencing of amplicons from four additional *V. alfalfae* isolates was carried out. As with *VnaChtBP*, no allelic polymorphisms were found among the sequences obtained and comparison of *V. nonalfalfae* and *V. alfalfae* gene sequences from these isolates also showed 97% sequence identity. Within the 36 single nucleotide polymorphisms identified, only resulted in 13 amino acid substitutions (Suppl. File S2).

### VnaChtBP forms dimers and has two potential binding sites for interaction with chitin

*V. nonalfalfae VnaChtBP* is an intronless gene and predicted to encode for a cysteine rich (12.5%) apoplastic effector (VnaChtBP) with 400 amino acids, including N-terminal signal peptide and six type 1 Chitin binding domains (ChtBD1; PF00187). This domain is classified in the CAZY database (Lombard et al. 2014) as Carbohydrate-binding module 18 (CBM18) and consists of 30 to 43 residues rich in glycines and cysteines, which are organized in a conserved four-disulfide core (Wright et al. 1991; Andersen et al. 1993; Asensio et al. 2000). It is a common structural motif, with a consensus sequence X3CGX7CX4CCSX2GXCGX5CX3CX3CX2 (Prosite PS50941), found in various plant and fungal defense proteins and is involved in the recognition and/or binding of chitin subunits (Finn et al. 2014).

Since many chitin binding proteins have been reported to form dimers (Liu et al. 2012; Sánchez-Vallet et al. 2013; Cao et al. 2014), a yeast-two-hybrid assay was carried out using *VnaChtBP* both as bait and prey to study the ability to dimerize. Dimer formation of VnaChtBP was detected on a minimal media using histidine as a reporter (Fig. 3A). Consistent with a weak interaction, only limited growth was observed on triple dropout reporter media (synthetic complete medium without leucine, tryptophan and uracil) and the X-gal reporter was not activated.

**Fig. 3.**
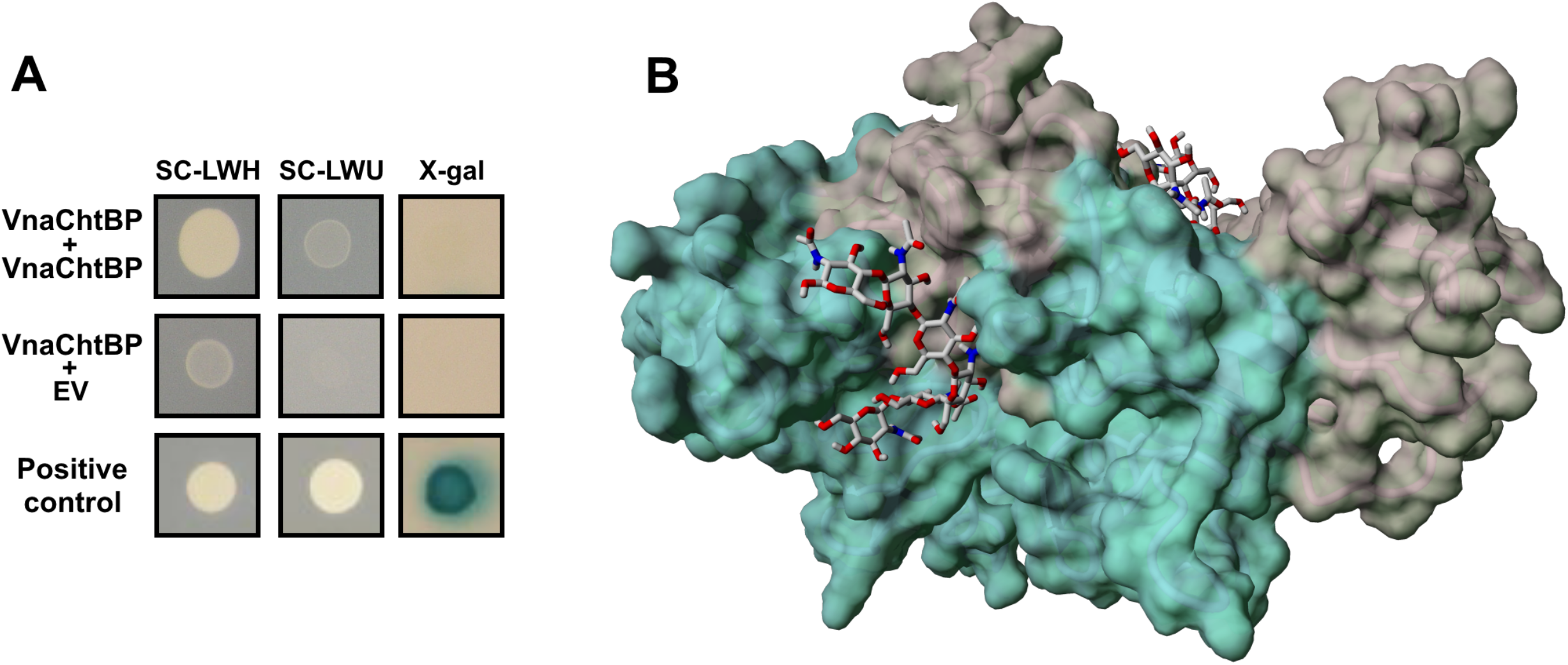
Confirmation of VnaChtBP dimerization (A) and schematic representation of the VnaChtBP homology model in complex with chitin hexamer (B). A: The effector gene *VnaChtBP* was cloned into the vectors pDEST22 and pDEST32 to serve both as bait and as prey and yeast-two-hybrid assay was performed. Weak dimerization of the effector was confirmed on a triple dropout reporter media SC-LWH and no self-activation of the pDEST22 construct with empty pDEST32 vector was detected on the X-gal reporter. B: The 3D model of VnaChtBP obtained by Swiss-Model (Arnold et al. 2006; Waterhouse et al. 2018) was refined by YASARA Structure (Krieger and Vriend 2014, 2015) and used in YASARA’s AutoDock VINA module (Trott and Olson 2009) for molecular docking of chitin hexamer, built in the SWEET PROGRAM (Bohne et al. 1998, 1999). VnaChtBP is in dimeric form, the chitin binding domains of the Chain A (Chain B) are in cyan (grey) color shades. The chitin hexamer is shown in stick representation.

To understand the chitin binding mechanism of CBM18 effectors better, homology modelling of VnaChtBP 3D structure was performed. The SWISS-MODEL server produced three models based on different templates, shown in Table 1. Model02 provided the best fit for four out of six CBM18 modules and was used as the basis of the characterization. Molecular docking of the chitin hexamer into the VnaChtBP model (Fig. 3B) shows that each protein monomer contributes to the formation of two binding sites accessible to the ligand. In binding site I (BSI), chitin hexamer is accommodated in a shallow groove formed by four hevein domains of polypetide chain A and two hevein domains of chain B, while binding site II (BSII) is comprised of four hevein domains of chain B and two domains of chain A. According to the analysis of the presented complex with YASARA, the binding of chitin hexamer in the BSI is strengthened by eleven (four accepted and seven donated) hydrogen bonds and eight hydrophobic interactions, which contribute to the total binding energy of 6.891 kcal/mol (AutoDock/Vina) and an estimated dissociation constant of 8.88 μM. Similar preference for binding of chitin hexamer in the BSII was observed, with an estimated dissociation constant of 2.01 μM and the total binding energy of 7.772 kcal/mol, supported by eight (three accepted, five donated) hydrogen bonds and 12 hydrophobic interactions between the ligand and receptor.

**Table 1.** The results of homology modelling of the VnaChtBP using the SWISS-MODEL server.

|  | Template | Description | Seq Identity | Oligo-States | GMOE | QMEAN | Coverage |
| --- | --- | --- | --- | --- | --- | --- | --- |
| Model01 | 2uwg.1.A | Wheat germ lectin | 43.98% | Homo-dimer | 0.31 | -4.61 | 98-318 |
| Model02 | 2wgc.1.A | Agglutin isolectin 1 | 40.96% | Homo-dimer | 0.32 | -3.22 | 39-254 |
| Model03 | 1ulk.1.A | Lectin-C | 49.15% | Homo-dimer | 0.20 | -0.65 | 212-378 |

### VnaChtBP binds chitin *in vitro* and protects fungal hyphae against plant chitinases

To confirm carbohydrate binding, *E. coli* produced and Ni-NTA affinity purified recombinant VnaChtBP (Suppl. Fig. S1) was used in a sedimentation assay with various carbohydrates. VnaChtBP binds specifically to chitin polymer, in the form of chitin beads and crab shell chitin, but not to the plant cell wall polymers cellulose and xylan (Fig. 4). To examine the affinity of VnaChtBP binding to chitin in more detail, recombinant protein was immobilized to the CM5 sensor chip and the VnaChtBP interaction with chitin hexamer was analyzed using surface plasmon resonance (SPR) (Kastritis and Bonvin 2013). VnaChtBP reveals concentration-dependent binding of chitin hexamer (Fig. 5) with a dissociation constant of 0.78 ± 0.58 μM, while no specific binding to other tested carbohydrates was detected (Suppl. Fig. S2). Since the chitin binding affinity of the protein increases for longer chitin oligomers (Asensio et al. 2000), this value is comparable to other reported chitin oligomer binding affinities of fungal effectors but exceeds by one order of magnitude those reported for *Arabidopsis* chitin recognition receptors and hevein (Table 2).

**Fig. 4.**
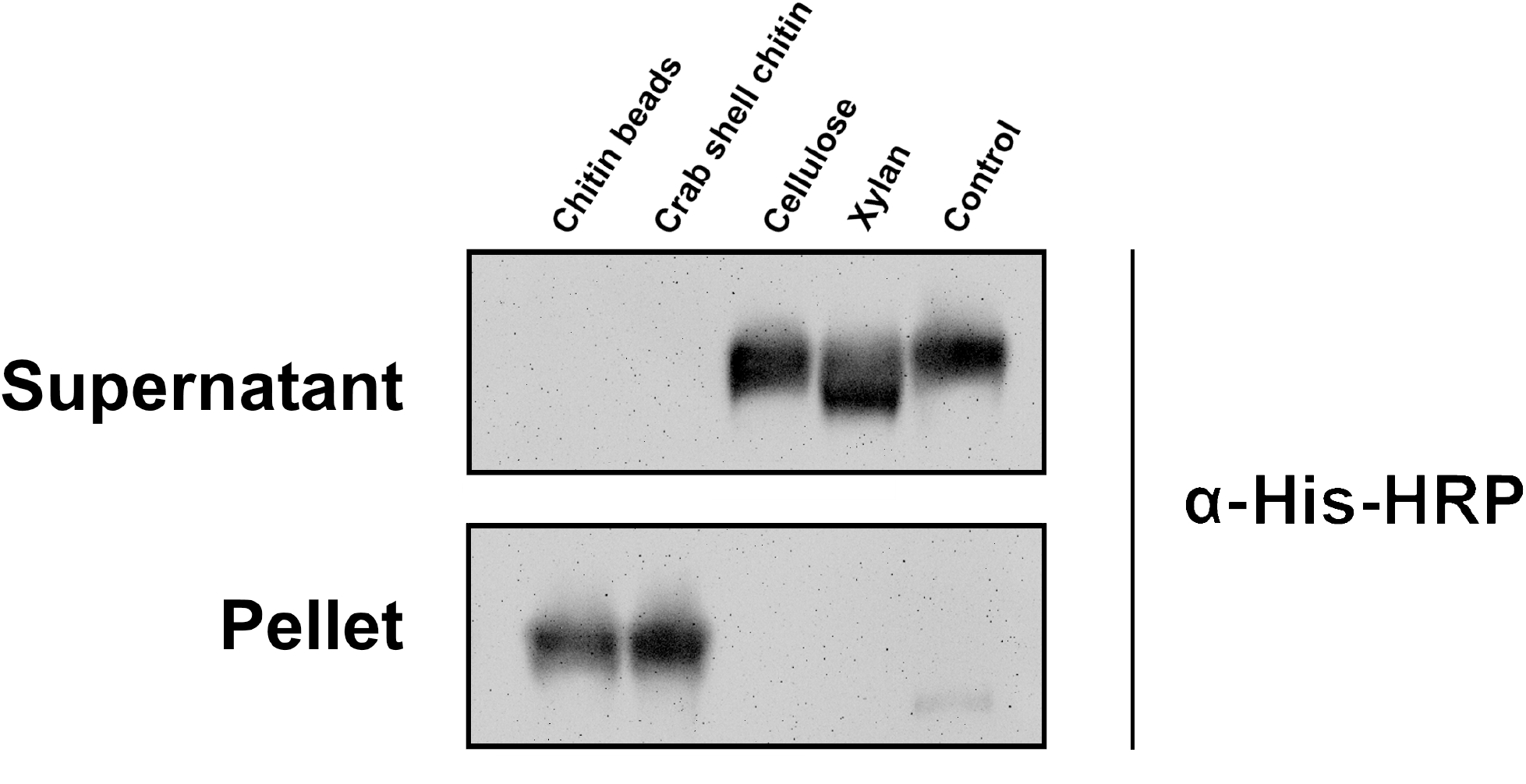
A carbohydrate sedimentation test confirmed that the recombinant protein VnaChtBP specifically binds to chitin. A recombinant protein (15 μg) that bound to chitin beads and crab shell chitin was detected in the sediment, while it was present in the supernatant when incubated with cellulose, xylan or without the addition of carbohydrates (control). Western blot analysis was performed with primary antibody His-probe (H-3) (SCBT) (1:1,000) and secondary Chicken anti-mouse IgG-HRP (SCBT) (1:5,000). Protein bands were detected using Super Signal West Pico (ThermoFisher Scientific) ECL substrate in a GelDoc-It2 Imager (UVP).

**Fig. 5.**
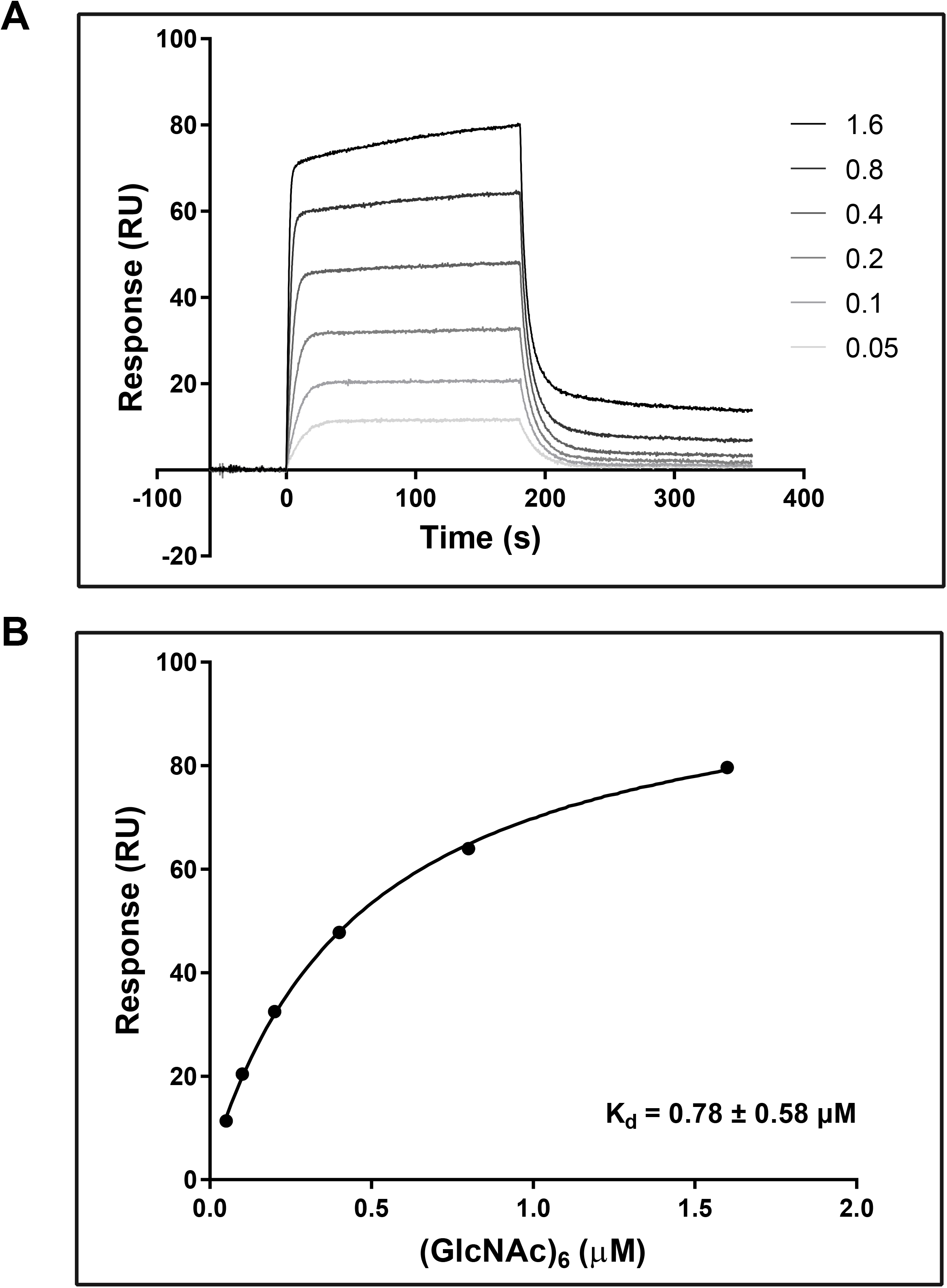
SPR analysis of chitin hexamer binding to VnaChtBP. Different concentrations (0.05, 0.1, 0.2, 0.4, 0.8, 1.6, 3.2 and 6.4 μM) of (GlcNAc)_6_ were tested for the binding (top panel). The binding curve (bottom panel) was generated by fitting steady state response levels at the end of the association phase, versus the concentration of the injected chitin hexamer. K_d_ was obtained by fitting the data to the steady-state affinity model. For reproducibility of binding, three independent titration experiments were performed. (GlcNAc)_6_, hexa-N-acetyl chitohexaose

**Table 2.** Comparison of chitin oligomer binding affinities of various fungal effectors and plant defense proteins, obtained using ITC or SPR.

| Organism | Protein | CAZy | Ligand | K<sub>d</sub> (μM) | Method | Reference |
| --- | --- | --- | --- | --- | --- | --- |
| *Verticillium nonalfalfae* | VnaChtBP | CBM18 | (GlcNAc)<sub>6</sub> | 0.78 ± 0.58 | SPR | This study |
| *Cladosporium fulvum* | Avr4 | CBM14 | (GlcNAc)<sub>6</sub> | 6.3 ± 0.23 | ITC | (van den Burg et al. 2004) |
| *Cladosporium fulvum* | Ecp6 | CBM50 | (GlcNAc)<sub>4,5,6,8</sub> | 11.5 to 3.7 | ITC | (de Jonge et al. 2010) |
| *Cladosporium fulvum* | Ecp6 | CBM50 | (GlcNAc)<sub>8</sub> | 1.3 × 10<sup>-3</sup> | SPR | (Mentlak et al. 2012) |
| *Magnaporthe oryzae* | Slp1 | CBM50 | (GlcNAc)<sub>8</sub> | 2.4 × 10<sup>-3</sup> | SPR | (Mentlak et al. 2012) |
| *Colletotrichum higginsianum* | ChELP1 | CBM50 | (GlcNAc)<sub>8</sub>-Bio | 2.6 × 10<sup>-5</sup> | SPR | (Takahara et al. 2016) |
| *Colletotrichum higginsianum* | ChELP2 | CBM50 | (GlcNAc)<sub>8</sub>-Bio | 2.5 × 10<sup>-4</sup> | SPR | (Takahara et al. 2016) |
| *Arabidopsis thaliana* | AtLYK5 | CBM50 | (GlcNAc)<sub>8</sub> | 1.72 | ITC | (Cao et al. 2014) |
| *Arabidopsis thaliana* | AtCERK1 | CBM50 | (GlcNAc)<sub>8</sub> | 455 | ITC | (Cao et al. 2014) |
| *Arabidopsis thaliana* | AtCERK1 | CBM50 | (GlcNAc)<sub>8</sub> | 448 | ITC | (Liu et al. 2012) |
| *Hevea brasiliensis* | Hevein | CBM18 | (GlcNAc)<sub>5</sub> | 2.1 | ITC | (Asensio et al. 2000) |

ITC, isothermal titration calorimetry; SPR, surface plasmon resonance, CBM, carbohydrate-binding module; GlcNAc, N-Acetylglucosamin, Bio, biotinylated

Chitin binding effectors have been reported to protect fungal hyphae from plant chitinases (van den Burg et al. 2004; Marshall et al. 2011). To determine whether recombinant VnaChtBP can protect fungal cell walls against hydrolysis by plant chitinases, a cell protection assay adapted from Mentlak et al. (2012) was performed using germinating conidia of *Trihoderma viride* and extracted xylem sap from *V. nonalfalfae* infected hop (Flajšman et al. 2017a) as a source of plant chitinases (containing 19 U of chitinase/mg total protein). In the presence of xylem sap, only minimal germination of the *T. viride* conidia occurred after 24 h incubation, while a pre-incubation in a 3 μM solution of recombinant VnaChtBP prior to the addition of xylem sap, enabled germination of conidia and hyphal growth. Interestingly, aggregation and compaction of fungal hyphae was detected only in the presence of both xylem sap and VnaChtBP, while normal mycelial growth without hyphal aggregation was observed in the solution of VnaChtBP (Fig. 6). We assume that VnaChtBP by binding and probably surrounding chitin fibers in the fungal cell wall, masks chitin and protects it from degradation by xylem sap chitinases.

**Fig 6.**
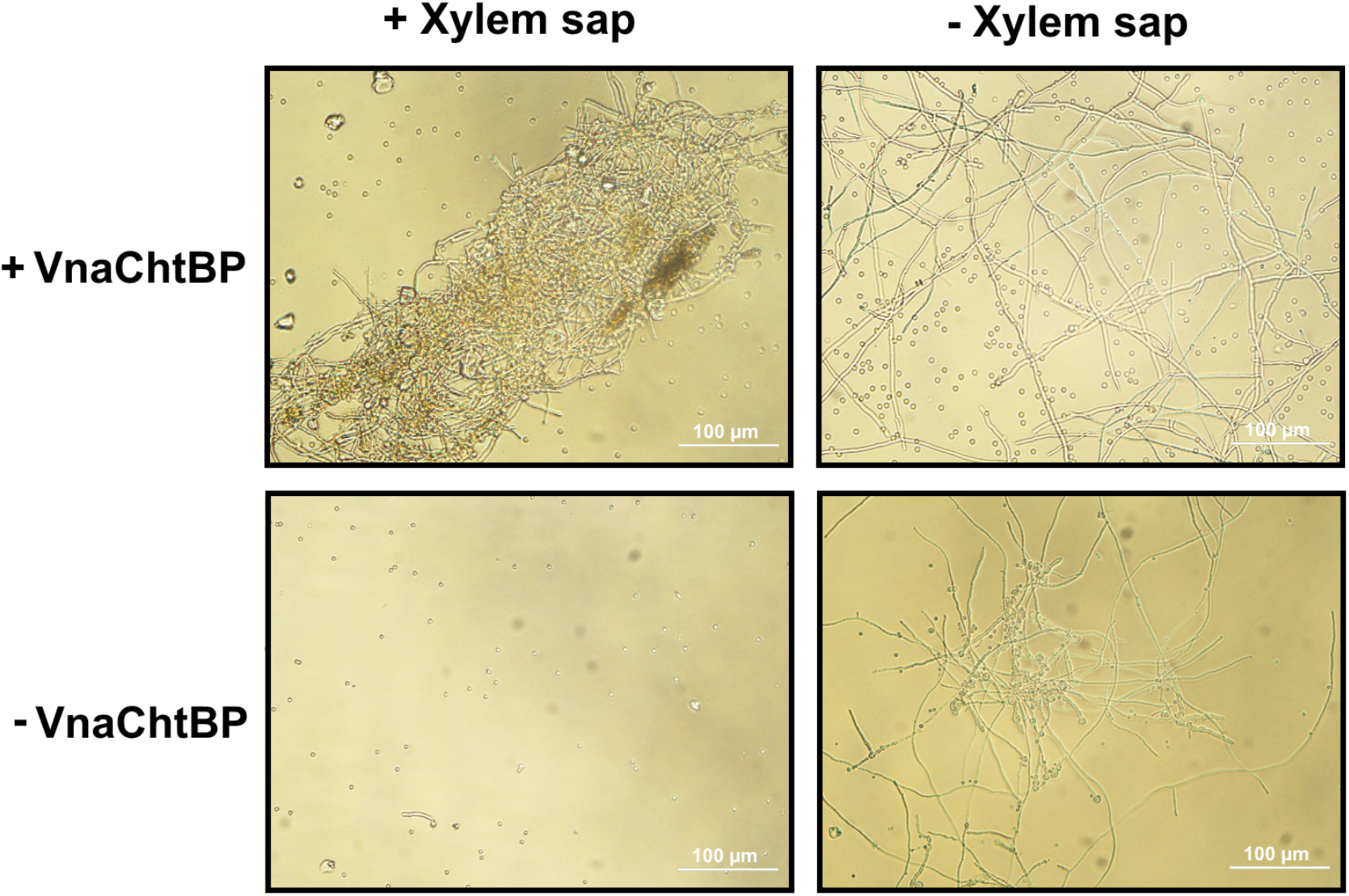
VnaChtBP protects fungus against degradation by plant chitinases. Micrographs of *Trihoderma viride* germinating spores, preincubated at RT for 2 h with 3 μM VnaChtBP, followed by the addition of xylem sap (19 U of chitinase/mg total protein) from *V. nonalfalfae* infected hop, were taken 24 h after treatment. The recombinant protein VnaChtBP caused aggregation and compaction of *T. viride* hyphae and protected the fungus from degradation by xylem sap chitinases. The chitinase activity of xylem sap was measured as a release of dye from Chitin Azure and one chitinase unit was defined as the amount of enzyme that caused a 0.01 increase in absorbance at 575 nm, measured at pH 5.0 and 25°C.

### *VnaChtBP* deletion has no significant effect on the growth and pathogenicity of *V. nonalfalfae*

Since *VnaChtBP* is specifically expressed during colonization of hop, its contribution to fungal virulence was tested in the susceptible hop variety ‘Celeia’. *V. nonalfalfae* knockout mutants of *VnaChtBP* were generated by targeted gene disruption via *A. tumefaciens*-mediated transformation. Prior to plant inoculation, growth of fungal colonies and sporulation of knockout mutants were assessed *in vitro* and compared to wild type. In the selected knockout mutants, mycelial growth and fungal morphology did not differ significantly from the wild type. Reduced sporulation was observed for both mutants compared to the wild type but this did not impact on disease frequency (Suppl. Fig. S3). After inoculation of hop plants, disease symptoms were independently assessed five times using a disease severity index (DSI) with a 0-5 scale (Radišek et al. 2003). After the final symptom assessment, the presence of fungus in all inoculated plants was confirmed through re-isolation tests and qPCR with fungus specific markers (Cregeen et al. 2015). Figure 7 shows symptom development in susceptible hop following infection with the wild type *V. nonalfalfae* and knockout mutants of *VnaChtBP*. Both *VnaChtBP* deletion mutants displayed Verticillium wilting symptoms (chlorosis and necrosis of the leaves) in susceptible hop similar to the wild type fungus, with no significant differences among them according to the DSI assessment. Independent pathogenicity assays with additional *VnaChtBP* deletion mutants yielded the same results (data not shown). This suggests that VnaCthBP function is redundant for *V. nonalfalfae* infection.

**Fig. 7.**
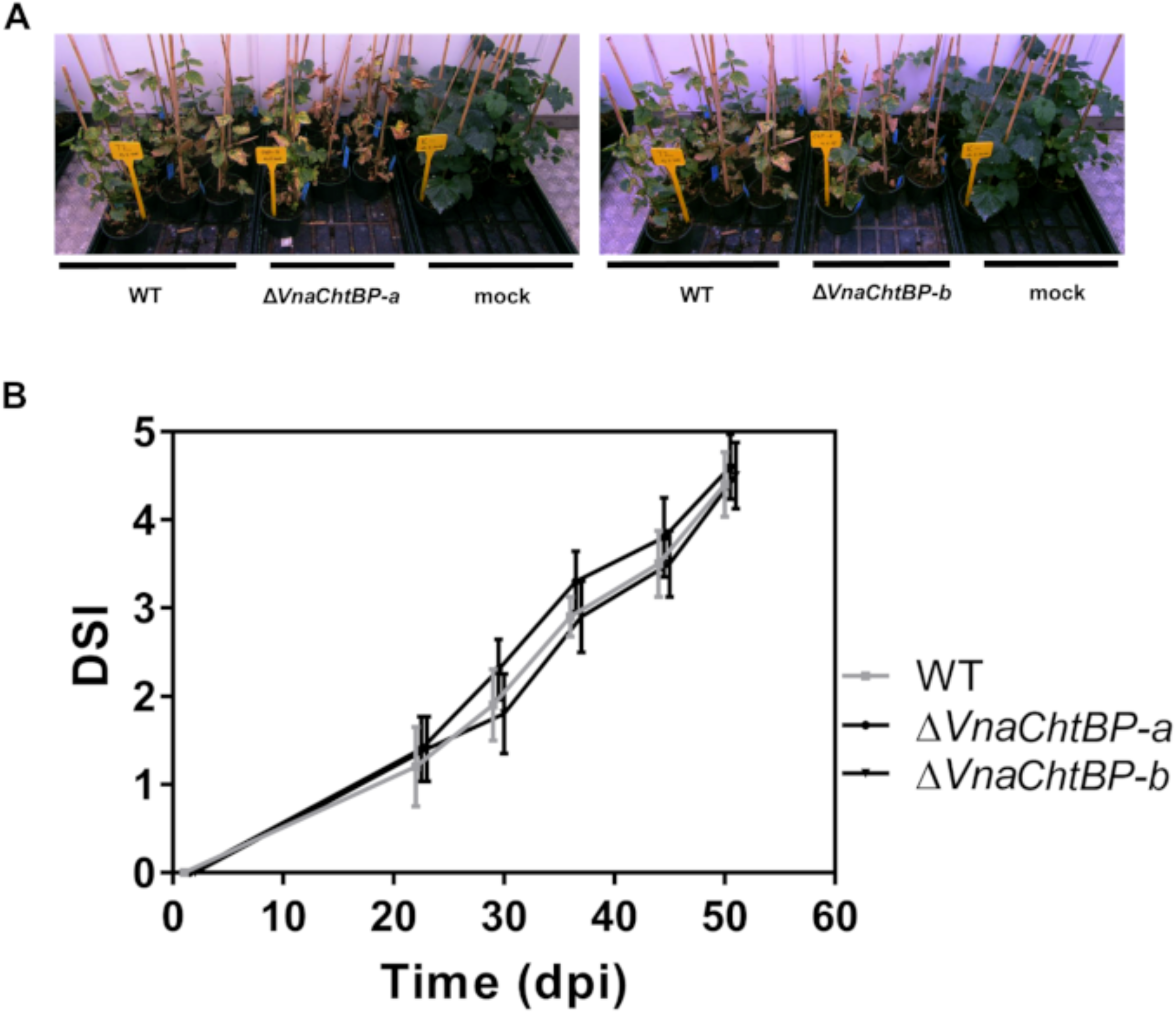
Symptom development in susceptible hop following infection with the wild type *V. nonalfalfae* and two knockout mutants of *VnaChtBP.* Plants of susceptible hop ‘Celeia’ were inoculated by root dipping in 5×10^6^ conidia/ml suspension and Verticillium wilting symptoms were assessed five times post inoculation. A: Both *VnaChtBP* deletion mutants displayed Verticillium wilting symptoms (chlorosis and necrosis of the leaves) in susceptible hop similar to the wild type fungus. Pictures were taken 35 days post inoculation. B: According to disease severity index (DSI) assessment with a 0-5 scale (Radišek et al. 2003) there were no significant differences between the wild type *V. nonalfalfae* and knockout mutants of *VnaChtBP.* Means with SE were calculated for 10 plants per treatment. Dpi, days post inoculation.

## Discussion

*V. nonalfalfae*, a soil born fungal pathogen, causes serious economic damage in European hop growing regions. Significant efforts have been invested to study the molecular mechanisms of Verticillium wilt in hop and fungus pathogenicity (Radišek et al. 2006; Jakše et al. 2013; Cregeen et al. 2015; Mandelc and Javornik 2015; Flajšman et al. 2016; Jakše et al. 2018; Marton et al. 2018).

*In planta* expressed fungal proteins are potential effector candidates, which might be implicated in pathogen virulence. The here studied effector candidate *V. nonalfalfae VnaChtBP*, encodes for a CBM18 domain containing chitin binding protein and is highly expressed in hop plants. Using an established bioinformatic pipeline (Marton et al. 2018), we identified eleven genes in the *V. nonalfalfae* genome that contain CBM18 domains. Of these genes, two harboured a single CBM18 domain (Fig. 1) and five, including *VnaChtBP*, contain a predicted N-terminal signal peptide. Although CBMs play a key role in the recognition of carbohydrates and are known to promote efficient substrate hydrolysis as a part of carbohydrate-active enzymes (e.g., CBM18 motifs found in chitinases), they have also been found to be present in toxins, virulence factors or pathogenesis-associated proteins (Guillén et al. 2010). Proteins containing CBM18 motifs are common in fungi, particularly in plant and animal pathogens. Indeed, they are almost three times as common in the proteomes of pathogens than in those of non-pathogenic fungi across the phylum Ascomycota (Soanes et al. 2008). Intriguingly, in *Verticillium spp*., CBM18 containing genes are more frequently observed in saprophytic *V. tricorpus* (13) (Seidl et al. 2015) than in pathogenic *V. dahliae* and *V. alfalfae* (9). The expansion of CBM18 domains in ChtBPs, could be linked to the evolution of pathogenicity, and has, for example, been reported in the fungal pathogen *B. dendrobatidis*, which has caused a worldwide decline of the amphibian populations (Abramyan and Stajich 2012). In total, eighteen genes with between one to eleven CBM18 domains have been identified in *B. dendrobatidis* including some classified as lectin-like proteins. Biochemical characterization of three such lectin-like proteins revealed that two have a signal peptide and co-localize with chitinous cell wall in *Saccharomyces cerevisiae*. Furthermore, one of these proteins has been shown to bind chitin and thereby protect *Trichoderma reesei* from exogenous chitinase, suggesting a role of lectin-like proteins in fungal defence (Liu and Stajich 2015). Similarly, in the rice blast fungus *M. oryzae*, 15 genes with one to four CBM18 domains were found, although gene-targeted disruption and tolerance to chitinase treatment did not support the implication of the tested genes in fungal pathogenicity (Mochizuki et al. 2011).

VnaChtBP consists of six tandemly repeated CBM18 motifs, contains a signal peptide and is predicted to reside in the apoplasm, which is consistent with the role of chitin binding in the extracellular space. Homology search of proteins that contain CBM18 motifs in other *Verticillium* species, revealed that this type of protein is common in pathogenic *Verticillium* species but it seems not to be ubiquitous. For example, in the recently sequenced genomes of five *V. dahliae* strains isolated from strawberry, three strains harbored ChtBPs with five, six and ten CBM18 motifs, while none were detected in two other strains.

Monitoring the *in planta* expression of *VnaChtBP* showed that it is highly expressed at the later stages of infection in a susceptible hop cultivar, and continuous to be expressed even at 30 dpi, when plants exhibit severe wilting symptoms (Cregeen et al. 2015; Marton et al. 2018). In contrast, in a resistant cultivar, the *VnaChtBP* gene is slightly induced after infection and then completely down-regulated. The expression pattern of the *VnaChtBP* gene coincides with *V. nonalfalfae* colonization of hop, whereby the fungus spread is unimpeded in susceptible plants while colonization is arrested around 12-20 dpi in resistant hop plants, presumably due to strong plant resistance responses (Cregeen et al. 2015). The immune reaction in incompatible interaction is unlikely to impose selection pressure on the *VnaChtBP* gene since no allelic polymorphisms were detected among analysed *V. nonalfalfae* isolates. Similarly, no allelic variation was found in the closest (97% identity) homolog to the *VnaChtBP* gene from isolates of *V. alfalfae*, suggesting highly conserved genes. Allelic variation is commonly detected in fungal proteins that function as avirulent (Avr) determinants upon perception by the host defence, but not necessarily in virulence factors of the pathogen (Stergiopoulos et al. 2007). Taken together, we speculate that the absence of allelic variation and the high gene expression observed *in planta* suggest a role for VnaChtBP in virulence of *V. nonalfalfae*. However, in pathogenicity assay, *VnaChtBP* targeted deletion mutants were not significantly impaired in their hops infectivity compared to wild type fungus which is in line with functional redundancy. Unchanged virulence of deletion mutants, presumably due to functional redundancy, has been reported for two other tested CBM18-containing ChtBPs in M. oryzae (Mochizuki et al. 2011) and also for LysM fungal effector, Mg1LysM of *Mycosphaerella graminicola* (Marshall et al. 2011). Indeed, other putative ChtBPs have been found in the *V. nonalfalfae* genome, which may have a role in protection of the fungal cell wall chitin or may interfere with chitin-triggered plant immunity. Specifically, Vna9.508, with one LysM domain, and Vna8.102, with five LysM domains, are both expressed during infection of hop and predicted to be secreted proteins with apoplastic localization (Marton et al. 2018). Orthologues of *C. fulvum* Avr4 with CBM14 chitin-binding motif were not identified in the *V. nonalfalfae* genome (Jakše et al. 2018) or in the predicted proteomes of other *Verticillium* species (Seidl et al. 2015).

Consistent with previously characterized CBM18 containing proteins from *M. oryzae* (Mochizuki et al. 2011) and *B. dendrobatidis* (Liu and Stajich 2015), recombinant VnaChtBP binds specifically to chitin beads and crab shell chitin, but not to plant cell wall cellulose or xylan. In addition to chitin polymer, recombinant VnaChtBP also binds chitin hexamer in an SPR experiment, with binding affinity in the submicromolar range. Compared to plant chitin receptors, recombinant VnaChtBP alongside LysM effectors Ecp6 from *C. fulvum*, Slp1 from *M. oryzae* (Mentlak et al. 2012), ChELP1 and ChELP2 from *C. higginsianum* (Takahara et al. 2016) exhibit three to five orders of magnitude higher affinity to chitin oligomers. It is thus not surprising that these fungal effectors are able to outcompete plant chitin receptors, such as *Arabidopsis thaliana* AtLYK5 (Cao et al. 2014) and AtCERK1 (Liu et al. 2012).

Based on NMR studies and solved crystal structures of plant LysM chitin receptors, several mechanisms for binding of chitin have been proposed; from a simple ‘continuous groove’ model for AtCERK1 (Liu et al. 2012) to the OsCEBiP ‘sandwich’ (Hayafune et al. 2014) and ‘sliding mode’ model (Liu et al. 2016). However, these models have been unable to explain the observed elicitor activities of chitin oligomers. Building on these models and using a range of chitosan polymers and oligomers bound to *Atcerk1* mutants resulted in an improved ‘slipped sandwich’ model that fits all experimental results (Gubaeva et al. 2018). A recent structural study of fungal LysM effector Ecp6 from *C. fulvum* revealed a novel chitin binding mechanism that explained how LysM effectors can outcompete plant host receptors for chitin binding (Sánchez-Vallet et al. 2013). Ecp6 consists of three tightly packed LysM domains, with a typical βααβ fold. Intra-chain dimerization of chitin-binding regions of LysM1 and LysM3 leads to the formation of a deeply buried chitin binding groove with an ultra-high (pM) affinity. The remaining LysM2 domain also binds chitin, albeit with low micromolar affinity, and interferes with chitin-triggered immunity, possibly by preventing chitin immune receptor dimerization and not by chitin fragment sequestering, as in case of LysM1-LysM3.

To date, to the best of our knowledge, the molecular mechanism of chitin binding of CBM18 fungal effectors remains elusive. However, the 3D homology model of VnaChtBP provides a tangible model for the molecular docking of the chitin hexamer. Although only four out of six CBM18 domains could be reliably modelled, the analysis revealed that VnaChtBP dimerizes. Importantly, this prediction was independently validated through a yeast-two-hybrid experiment. The VnaChtBP complex has two putative chitin binding sites which form a shallow binding cleft by cooperation of both polypeptide chains and have a similar preference to chitin. As in CBM18 lectin-like plant defence proteins (Jiménez-Barbero et al. 2006), typically represented by a small antifungal protein hevein from the rubber tree (*Hevea brasiliensis*), a network of hydrogen bonds and several hydrophobic interactions occur between VnaChtBP residues and N-acetyl moieties of the chitin oligomer. These are thought to stabilize the interaction and contribute to submicromolar chitin binding affinity, as determined by SPR. Similarly, the recently solved crystal structure of fungal effector CfAvr4, a CBM14 lectin, in complex with chitin hexamer (Hurlburt et al. 2018) has revealed that two effector molecules form a sandwich structure, which encloses two parallel stacked chitin hexamer molecules, shifted by one sugar ring, in an extended chitin binding site. In this complex, the interaction is mediated through aromatic residues and numerous hydrogen bonds with both side chains and main chains. Interestingly, no intermolecular protein-protein interactions have been observed across the dimer, suggesting ligand induced effector dimerization. Site-directed mutagenesis of residues responsible for binding of chitin hexamer showed that ligand binding function is independent from recognition by host resistance protein Cf-4.

Fungal plant pathogens have evolved several strategies to escape the surveillance of chitin-related immune systems (Sanchez-Vallet et al. 2014). The different mechanisms used include conversion of chitin to chitosan by chitin deacetylases and inclusion of α-1,3-glucan in the cell walls, as well as secretion of diverse effectors that can shield the fungal hyphae from hydrolysis by plant chitinases, directly inhibiting their activity, acting as scavengers of chitin fragments or preventing chitin–induced receptor dimerization. Secreted effector Avr4 from *C. fulvum* binds to fungal cell wall chitin to reduce its accessibility to host chitinases (van den Burg et al. 2006). Similar to CfAvr4, wheat pathogen *M. graminicola* secreted effectors Mg1LysM and Mg3LysM protect fungal hyphae from hydrolysis by plant chitinases (Marshall et al. 2011). We provide evidence that, in addition to Avr4 (CBM14) and LysM (CBM50) effectors, structurally unrelated CBM18 lectin-like proteins that are found in fungal pathogens of plants (this study) and amphibian pathogens (Liu and Stajich 2015) have evolved a chitin shielding ability against plant chitinases.

## Materials and Methods

### Maintenance of plant cultures and cultivation of microorganisms

*Nicotiana benthamiana* seedlings were grown at 23 ± 2°C under a 16:8 photoperiod. Hop (*Humulus lupulus* L.) of susceptible ‘Celeia’ and resistant ‘Wye Target’ cultivars were grown as described previously (Flajšman et al. 2017b). *Escherichia coli* MAX Efficiency DH5α or MAX Efficiency DH10B (both from Invitrogen, ThermoFisher Scientific) were used for plasmid propagation and were grown at 37°C on LB agar plates or liquid medium supplemented with appropriate antibiotics (carbenicillin 100 mg/liter, kanamycin 50 mg/liter, spectinomycin 100 mg/liter or gentamicin 25 mg/liter). *E. coli* Shuffle T7 (New England Bioloabs) were propagated at 30°C and protein expression was performed at 16°C. *Trichoderma viride* was obtained from The Microbial Culture Collection Ex (IC Mycosmo (MRIC UL)) and all *Verticillium* strains were from the Slovenian Institute of Hop Research and Brewing fungal collection. Fungi were grown at 24°C in the dark on ½ Czapek-Dox agar plates or liquid medium. Knockout mutants were retrieved from selection medium supplemented with 150 mg/liter timentin and 75 mg/liter hygromycin. For agroinfiltration, *Agrobacterium tumefaciens* MV3101 was grown at 28°C on YEB agar plates or liquid medium supplemented with rifampicin 25 mg/liter, gentamicin 25 mg/liter, and spectinomycin 50 mg/liter.

### RNA sequencing

RNA-Seq library preparation from *V. nonalfalfae* infected hop at 6, 12, 18 and 30 days post inoculation (dpi) and data processing have been previously described (Progar et al. 2017). Fungal transcripts were filtered out and their gene expression profiles were generated using the Hierarchical clustering with Euclidean distance method in R language (R Core Team 2016). Data were presented as a matrix of log2CPM (counts per million–number of reads mapped to a gene model per million reads mapped to the library) expression values.

### *VnaChtBP* gene expression profiling with RT-qPCR

The expression of *VnaChtBP* was quantified by RT-qPCR in hop infected with *V. nonalfalfae* isolate T2. Total RNA was extracted at 6, 12, and 18 dpi using a Spectrum Plant total RNA kit (Sigma-Aldrich) and 1 μg was reverse transcribed to cDNA using a High Capacity cDNA reverse transcription kit (Applied Biosystems). The qPCR reaction was run in 5 biological and 2 technical replicates on an ABI PRISM 7500 (Applied Biosystems), under the following conditions: denaturation at 95°C for 10 min, followed by 40 cycles at 95°C for 10 s, 60°C for 30 s, and consisted of: 50 ng of cDNA, 300 nM forward and reverse primer, and 5 μl of Fast SYBR Green master mix (Roche). The results were analyzed using the ΔΔCt method (Schmittgen and Livak 2008). Transcription levels of *VnaChtBP* were quantified relative to its expression in liquid Czapek-Dox medium and normalized to fungal biomass in hop using topoisomerase and splicing factor as reference genes (Marton et al. 2018). Primers used are listed in Table 3.

**Table 3.**
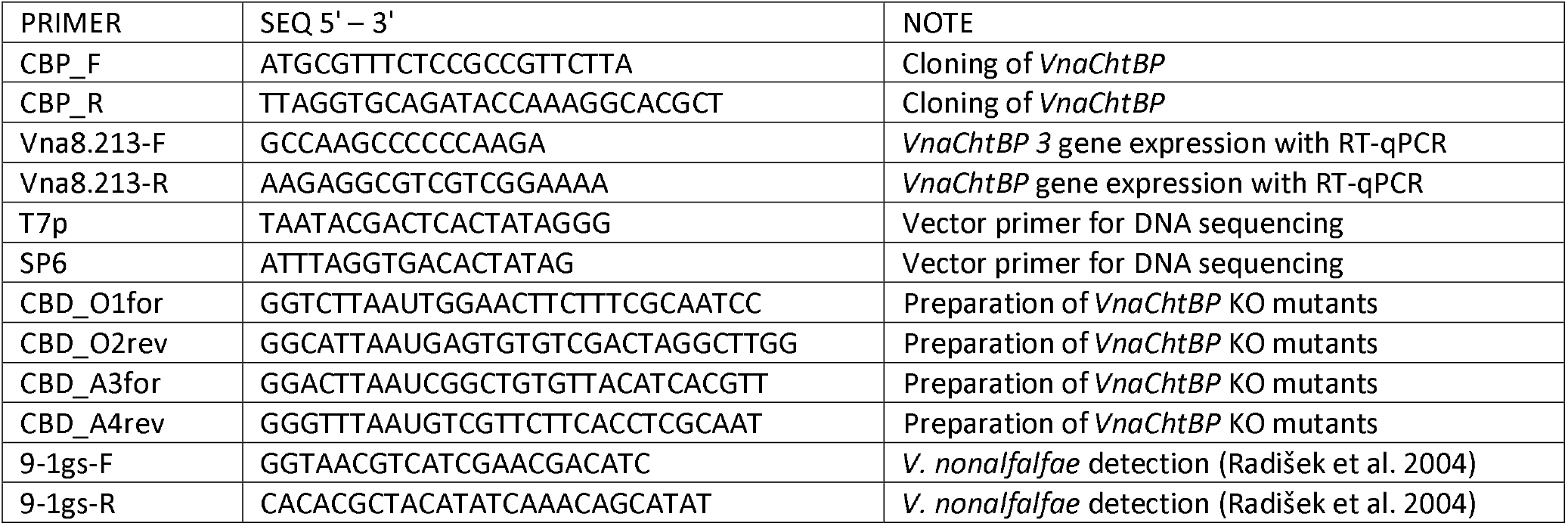
List of primers used for cloning, gene disruption and expression of VnaChtBP, and fungal identification.

### Genetic analysis

Genomic DNA was extracted from 7-10 day PDA cultured *Verticillium* isolates by the CTAB extraction method (Möller et al. 1992). PCR reactions were performed in 50 μl using Q5^®^ High-Fidelity DNA Polymerase (NEB), 500 nM gene-specific primers (Table 3) and 100 ng DNA under the following conditions: denaturation at 95°C for 10 min, followed by 40 cycles at 95°C for 10 s, 58°C for 30 s, 72°C for 90 s, and a final elongation step at 72°C for 90 s. PCR products were purified from agarose gel (Silica Bead DNA Gel Extraction Kit, Fermentas), cloned into pGEM®-T Easy vector (Promega) and sequenced using Sanger technology with gene-specific and plasmid-specific primers (Table 3). Sequences were analyzed using CodonCode Aligner V7.1.2 (CodonCode Co.) and deposited at the NCBI.

### Bioinformatic analysis

A putative localization of VnaChtBP to the apoplast was predicted with ApoplastP 1.0 (Sperschneider et al. 2017). To classify *V. nonalfalfae* CBM18-containing proteins functionally, sequence based searches were carried out using the FunFHMMer web server at the CATH-Gene3D database (Dawson et al. 2017). To obtain VnaChtBP homologs, the amino acid sequence of VnaChtBP was used as a query for NCBI BLAST+ against UniProt Knowledgebase at Interpro (Li et al. 2015).

### Yeast-two-hybrid assay

Dimerization of VnaChtBP was examined with a yeast-two-hybrid experiment using the ProQuest Y2H system (Invitrogen). To generate bait and prey vectors, the *VnaChtBP* gene was cloned into pDEST22 and pDEST32, respectively, and co-transformed in yeast. The interactors were confirmed by plating the yeast co-transformants on triple dropout reporter medium SC-LWH (synthetic complete medium without leucine, tryptophan and histidine), on triple dropout reporter medium SC-LWU (synthetic complete medium without leucine, tryptophan and uracil) and by performing an X-gal assay. The self-activation test of a pDEST22 construct containing *VnaChtBP* gene with empty pDEST32 vectors was also performed.

### 3D modelling and molecular docking

The SWISS-MODEL (Arnold et al. 2006; Waterhouse et al. 2018) server produced three models based on different templates and Model02 was selected for further modelling. The output protein structure was additionally minimized in explicit water using an AMBER14 force field (Duan et al. 2003) and ‘em_runclean.mcr’ script within YASARA Structure (Krieger and Vriend 2014, 2015). A 3D structure model of chitin hexamer was built with SWEET PROGRAM v.2 (Bohne et al. 1998, 1999), saved as a PDB file and used as a ligand in subsequent molecular docking experiments with AUTODOCK/VINA (Trott and Olson 2009), which is incorporated in YASARA Structure. To ensure the integrity of docking results, 200 independent dockings of the ligand to the receptor were performed. The pose with the best docking score was selected for further refinement using ‘md_refine.mcr’ script provided by YASARA Structure. The final model of the hexameric chitin bound to the VnaChtBP dimer was then used for the analysis.

### Recombinant protein production

*VnaChtBP* DNA without predicted signal peptide (SignalP 4.1) was cloned into a pET32a expression vector using a Gibson Assembly^®^ Cloning Kit (NEB). The protein expression in *E. coli* SHuffle^®^ T7 cells (NEB) was induced at OD600 = 0.6 with 1 mM IPTG and incubated overnight at 16°C. The recombinant protein was solubilized from inclusion bodies using a mild solubilization method (Qi et al. 2015). Briefly, pelleted cells were resuspended in cold PBS buffer and disrupted by sonication. After centrifugation, the pellet was washed with PBS, resuspended in urea, frozen at −20°C and slowly allowed to thaw at RT. The recombinant protein was purified using Ni-NTA Spin Columns (Qiagen) according to the manufacturer’s protocol, aliquoted and stored in 20 mM Tris (pH 8.0) at −80°C.

### Carbohydrate sedimentation assay and Western blot detection

The carbohydrate sedimentation assay was adapted from (van den Burg et al. 2006). Briefly, 15 μg of recombinant VnaChtBP in 20 mM Tris (pH 8.0) was mixed with 1.5 mg of chitin magnetic beads (NEB), crab shell chitin (Sigma-Aldrich), cellulose (Sigma-Aldrich) or xylan (Apollo Scientific) and incubated at RT for 2 h on an orbital shaker at 350 rpm. The same amount of protein in Tris buffer without added carbohydrates was used as a negative control. After centrifugation (5 min, 13,000 g), the supernatant was collected and the pellet was washed three times with 800 μl 20 mM Tris (pH 8.0) prior to resuspension in 4X Bolt™ LDS Sample Buffer with the addition of reducing agent (Invitrogen).

The presence of VnaChtBP in different fractions was determined by WB analysis. Samples (25 μL) were loaded on a precast Bolt™ 4-12% Bis-Tris gel (Invitrogen) and SDS-PAGE in 1x MOPS running buffer was performed using a Mini Gel Tank (ThermoFisher Scientific) for 30 min at 200 V. Proteins were transferred for 1 h at 30 V to an Invitrolon PVDF membrane (Invitrogen) and Ponceau S stained. The membrane was blocked with 5% BSA in 1x PBS before the addition of the primary antibody His-probe (H-3) (SCBT) (1:1,000). The membrane was incubated overnight at 4°C, washed with 1x PBS and incubated in a solution of secondary Chicken anti-mouse IgG-HRP (SCBT) (1:5,000) for 1 h. Protein bands were detected using Super Signal West Pico (ThermoFisher Scientific) ECL substrate in a GelDoc-It2 Imager (UVP).

### Surface plasmon resonance

The binding of hexa-N-acetyl chitohexaose ((GlcNAc)_6_; IsoSep) to VnaChtBP was measured using a Biacore T100 analytical system and CM5 sensor chip (Biacore, GE Healthcare). The CM5 sensor chip was activated using an Amine coupling Kit (GE Healthcare) according to the manufacturer’s instructions. VnaChtBP was diluted into 10 mM Sodium acetate pH 5.1 to a final concentration of 0.1 mg/ml and injected for five minutes over the second flow cell. The first flow cell was empty and served as a reference cell to control the level of non-specific binding. The final immobilization level was approximately 10,000 response units (RU). The (GlcNAc)_6_ stock solution was diluted into a series of concentrations (0.05, 0.1, 0.2, 0.4, 0.8, 1.6, 3.2 and 6.4 μM) with the HBS buffer (10 mM HEPES, 140 mM NaCl, pH 7.4) and assayed to detect direct binding to VnaChtBP. Titration was performed in triplicate. In addition to chitin hexamer, N-acetyl glucosamine, glucosamine, glucose, galactose and mannose were tested at a 500 μM concentration in HBS buffer. Biacore T100 Evaluation software was used to assess the results. First, the sensorgrams were reference and blank subtracted, then a Steady State Affinity model was applied to calculate the affinity constant (K_d_). The average of three repeated experiments was used for final Kd determination.

### Xylem sap extraction and chitinase activity assay

Xylem sap was extracted from infected hop plants in a pressure chamber at 0.2 MPa for 120 min (Flajšman et al. 2017a). Chitinase activity of xylem sap was determined by mixing 150 μl of xylem sap, or 100 mM Na-acetate (pH 5.0) buffer as negative control, with 1.5 mg of Chitin Azure (Sigma-Aldrich) dissolved in 150 μl 100 mM Na-acetate (pH 5.0). The samples were incubated for 150 min at 25°C on a rotatory shaker at 70 rpm. An aliquot of 80 μl was taken immediately (blank sample) and after 150 min. The reaction was stopped with the addition of 20%v/v HCl and samples were centrifuged for 10 min at 10,000 g. The chitinase activity of xylem sap in the supernatant was determined by measuring the absorbance of released Remazol Brilliant Blue dye at 575 nm against 100 mM Na-acetate (pH 5.0). One enzyme unit was defined as the amount of chitinase that produced a 0.01 increase in absorbance at 575 nm, measured at 25°C and pH 5.0. The total protein concentration of the xylem sap was measured in a 10x diluted sample using a Pierce™ BCA Protein Assay Kit (Thermo Scientific) following the standard protocol.

### Cell Protection Assay

The cell protection assay was adapted from Mentlak et al., 2012. *Trihoderma viride* conidia were harvested, diluted to 2,000 conidia/ml in 50 μl ½ Czapek-dox medium and incubated overnight. After germination of the conidia, 25 μl of recombinant VnaChtBP (3 μM final concentration) or an equal volume of storage buffer (20 mM Tris; pH 8.0) were added and the conidial suspensions were incubated for 2 h. Fungal cell wall hydrolysis was triggered by the addition of 25 μl of xylem sap as a source of plant chitinases, while 25 μl of Na-acetate (100 mM; pH 5.0) was used in the control experiment. After 24 h incubation, mycelia formation and fungal growth were examined using a Nikon Eclipse 600 microscope.

### Pathogenicity assay

*VnaChtBP* knockout mutants were generated using the *Agrobacterium tumefaciens* mediated transformation protocol described previously (Flajšman et al. 2016) and primers listed in Table 3. Before pathogenicity tests were carried out, fungal growth and conidiation were inspected as described previously (Flajšman et al. 2017b). Ten plants of the Verticillium wilt susceptible hop cultivar ‘Celeia’ were inoculated by 10-min root dipping in a conidia suspension (5× 10^6^ conidia/ml) of two arbitrarily selected *VnaChtBP* knockout mutants. Conidia of the wild type *V. nonalfalfae* isolate T2 served as a positive control and sterile distilled water was used as a mock control. Re-potted plants were grown under control conditions in a growth chamber (Flajšman et al. 2017b). Verticillium wilting symptoms were assessed five times over seven weeks post-inoculation using a disease severity index (DSI) with a 0-5 scale (Radišek et al. 2003). After symptom assessment, a fungal re-isolation test (Flajšman et al. 2017b) and qPCR using *V. nonalfalfae* specific primers (Cregeen et al. 2015) were performed to confirm infection of the tested hop plants.

## Acknowledgments

This research was supported by the Slovenian Research Agency grants P4-0077, J4-8220 and fellowship 342257. This work benefitted from interactions promoted by the COST Action FA 1208 https://cost-sustain.org. The authors would like to thank Dr Vasja Progar for transcriptome analysis, Dr Vesna Hodnik for SPR analysis and Dr Miles Armstrong, Dr Marko Dolinar and Dr Jernej Jakše for technical advice.

